# Born to Condense: Polysomes Drive Co-Translational Condensation of Biomolecular Condensate Proteins

**DOI:** 10.1101/2025.09.30.679637

**Authors:** Zhouyi He, Chi Fung Willis Chow, Agnes Toth-Petroczy, Jens-Uwe Sommer, Tyler S. Harmon

**Affiliations:** Division Theory of Polymers, Leibniz Institute of Polymer Research, Dresden, 01069, Germany; Institute of Theoretical Physics, Technische Universität Dresden, Dresden, 01062, Germany; Max Planck Institute of Molecular Cell Biology and Genetics, Dresden, 01307, Germany; Center for Systems Biology Dresden, Dresden, 01307, Germany; Cluster of Excellence Physics of Life, Technische Universität Dresden, Dresden, 01062, Germany

**Keywords:** Biomolecular Condensates, Liquid-Liquid Phase Separation, Co-Translation Condensation, Polysome, Protein Domain Architecture

## Abstract

Biomolecular condensates formed by protein liquid-liquid phase separation (LLPS) are ubiquitous in cells and play crucial roles in cellular regulation. While the physics and functions of LLPS are well studied, its interplay with protein synthesis – translation – remains largely unexplored. Here we introduce a theoretical framework for Co-Translational Condensation (CTC), in which nascent protein chains of polysomes – multiple ribosomes on one mRNA – interact with condensates, localizing translation to condensate surfaces. Using coarse-grained simulations, we show that protein domain architecture dictates the thermodynamics of CTC, consistent with a Langmuir adsorption model. Bioinformatic analysis of more than 7,500 proteins reveals that most condensate-associated proteins have architectures favoring CTC, with strong interaction regions of nascent chains exposed on polysomes. At the dynamical level, simulation and reaction–diffusion modeling reveal that CTC is kinetically feasible within typical polysome lifetimes, either through large polysomes nucleating new condensates or via diffusion to pre-existing condensates. As a case study, we demonstrate that CTC enhances post-translational modifications by minimizing unmodified intermediates. More broadly, we anticipate CTC may also influence protein folding, misfolding, and signal-integration latency. Together, our results establish CTC as a general mechanism coupling translation with phase separation, with broad implications for protein evolution, cellular organization, and synthetic biology.

## 1 Introduction

Biomolecular condensates formed via liquid–liquid phase separation (LLPS) exemplify how living matter leverages soft condensed matter physics to organize its interior. These dynamic membrane-less compartments contribute to key functions such as RNA metabolism, stress granule assembly, and stress responses[1–4]. While the physics of LLPS itself[5] and its role in regulating processes like gene expression[2–4, 6] are increasingly well-understood, a fundamental question remains: how the spatiotemporal control of condensates is connected to its prerequisite – protein synthesis.

During translation, multiple ribosomes moving along an mRNA generate a bottlebrush-like polymer – the polysome –where the nascent chains act as densely grafted side chains (Fig. 1**a**). The number of ribosomes per mRNA (polysome size) depends on organism, transcript length, and translation rate, and has been observed to range from 1 to 40 [7–9]. This presents a rich physical problem: if the translated protein is a client or scaffold of a condensate, its own synthesis could localize to the condensate surface through interactions between the nascent chains and the condensed phase, a process we term Co-Translational Condensation (CTC) (Fig. 1**b**).

**Fig. 1:**
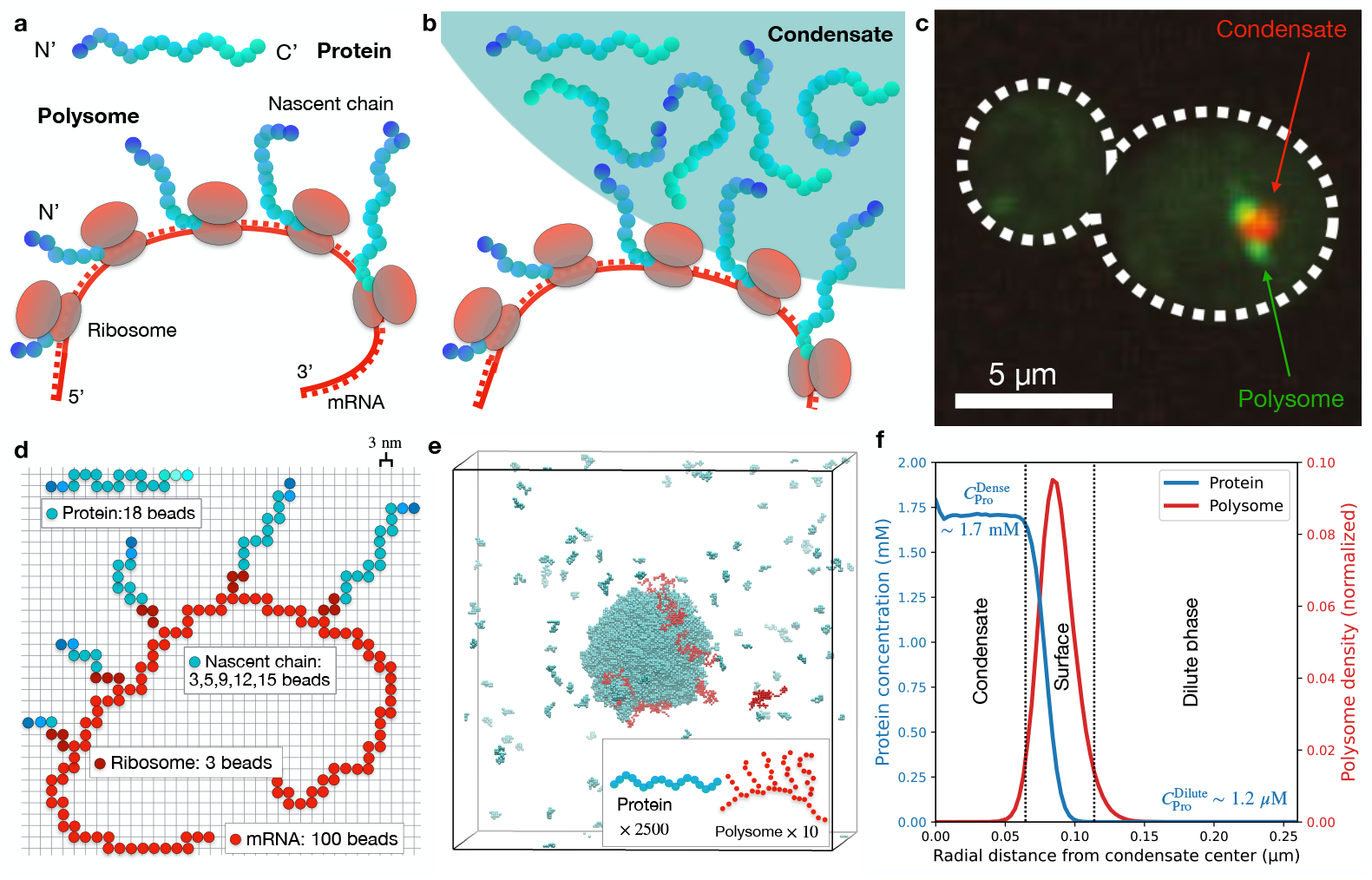
Co-Translational Condensation (CTC): polysomes and condensates of its encoded protein. **a**, Cartoon of a polysome that is translating condensate-associated proteins and the corresponding protein. Nascent protein chains thar are closer to the mRNA 3’ end are longer. The N-terminal of the proteins are at the tips of the nascent chains. **b**, Cartoon of a polysome’s nascent proteins interacting with a condensate. **c**, Experimental observation of CTC: polysomes (green) are co-localized with a condensate enriched in its encoded protein (red) in yeast, for more details on the experimental methodology, refer to Heidenreich *et al*. [10]. **d**, 3D lattice coarsegrained model of polysomes and proteins. Each lattice length measures 3 nm. A protein is coarse-grained as 18 beads, and polysomes are coarse-grained into small polysomes (5 nascent chains with 160 beads), small polysomes (9 nascent chains with 208 beads) or large polysomes (15 nascent chains with 265 beads), see SI Fig. 1 for detials. **e**, Representative simulation snapshot of 2500 proteins (homopolymers) and 10 polysomes in a cubic box with length of 600 nm. Most polysomes localize to condensate surfaces, see SI Movie II for the full trajectory. **f**, Radial distribution of material from the condensate center, time-averaged after system equilibration, quantifying the polysome localization. *c*_*Pro*_ denotes the concentration of protein.

Emerging experimental evidence supports this concept. For instance, polysomes in yeast localize to condensates of the proteins they encode [10] (Fig. 1**c**), and a similar mechanism facilitates the assembly of axonemal dynein complexes [11]. These observations point to a general physicochemical phenomenon driven by protein-protein interactions.

However, key physical questions remain open. What are the molecular determinants– the “sequence-to-phase” rules–that govern whether a polysome will associate with a condensate? What are the kinetic pathways for this association? And most importantly, what functional advantages does this coupling confer?

Here, we use multi-scale modeling to establish a biophysical foundation for CTC. We demonstrate that protein domain architecture dictates polysome-condensate affiliation, that CTC is a plausible pathway for most condensate-forming proteins, and that polysomes can associate via both diffusion and nucleation mechanisms. Furthermore, we show that CTC can enhance biochemical processes like post-translational modifications, illustrating a functional synergy between translation and phase separation. Our work suggests that CTC is a fundamental physical mechanism, shaped by evolution and ripe for exploration in synthetic biology, that functionally couples translation and condensation to control cellular organization.

## 2 Multi-scale modeling

We performed 3D lattice-based Monte Carlo polymer simulations[12], coarse-graining proteins into bead–spring polymers with excluded volume at a lattice length of ∼ 3 nm — corresponding to a folded domain or ∼30 disordered residues[13]. This approach enables simulations of 10^4^ polymers in 1 *µ*m^3^, and has been successfully used to study multiple condensates[14–23]; see Methods for details.

All proteins are represented as linear chains of 18 lattice beads (Fig. 1**d**). The corresponding polysome of each protein architecture is modeled as a branched polymer: the main chain represents the mRNA, and each branch represents a ribosome with its nascent protein chain (Fig. 1**d**). Because translation is directional, the N-terminal of each nascent chain is located at the branch tip. We considered two polysome sizes representing different ribosome packing density[7–9]: small polysomes with 5 nascent chains (160 beads per polysome), medium polysomes with 9 nascent chains (208 beads per polysome) and large polysomes with 15 nascent chains (265 beads per polysome), see Methods and SI.

In cells, polysomes typically disassemble after translating proteins for minutes to tens of minutes [24]. These timescales, and the corresponding cellular length scales, exceed the reach of coarse-grained simulations. Therefore, we used simulations to parameterize reaction–diffusion models that capture the emergent behavior at cellular scales.

## 3 Thermodynamics of Co-Translational Condensation

We begin by examining homopolymer-like proteins (Homo), in which all coarse-grained beads are identical and interact via uniform weak attractions. This model does not represent a literal homopolymer (e.g., poly-glycine), but rather an abstraction for proteins whose regions interact in a roughly equivalent manner[25], providing a baseline for understanding how architecture influences CTC. Fig. 1**e,f** and Supplementary Movie II show a representative simulation of Homo proteins forming a condensate, with a large fraction of their polysomes localizing translation at the condensate surface (78.7% *±* 0.5% CTC).

### 3.1 Protein architecture dictates Co-Translational Condensation

While the diversity of protein sequences is vast, we expect large-scale architectural features to control CTC, while small-scale variations are averaged-out. To capture these features, we used catGRANULE 2.0 [26], a tool that predicts LLPS propensity along the sequence, to study all condensate-associated proteins from the CD-CODE database[27]. The resulting LLPS propensity profiles motivated us to classify proteins into 5 distinct and biologically meaningful architectures to allow clear comparison based on the arrangement of “strong-interaction” regions (regions above average LLPS propensity); see Extended Data Fig.1 and Methods for classification details. Representative profiles[23, 28] of each architectures are shown in Fig. 2**a**:

**Fig. 2:**
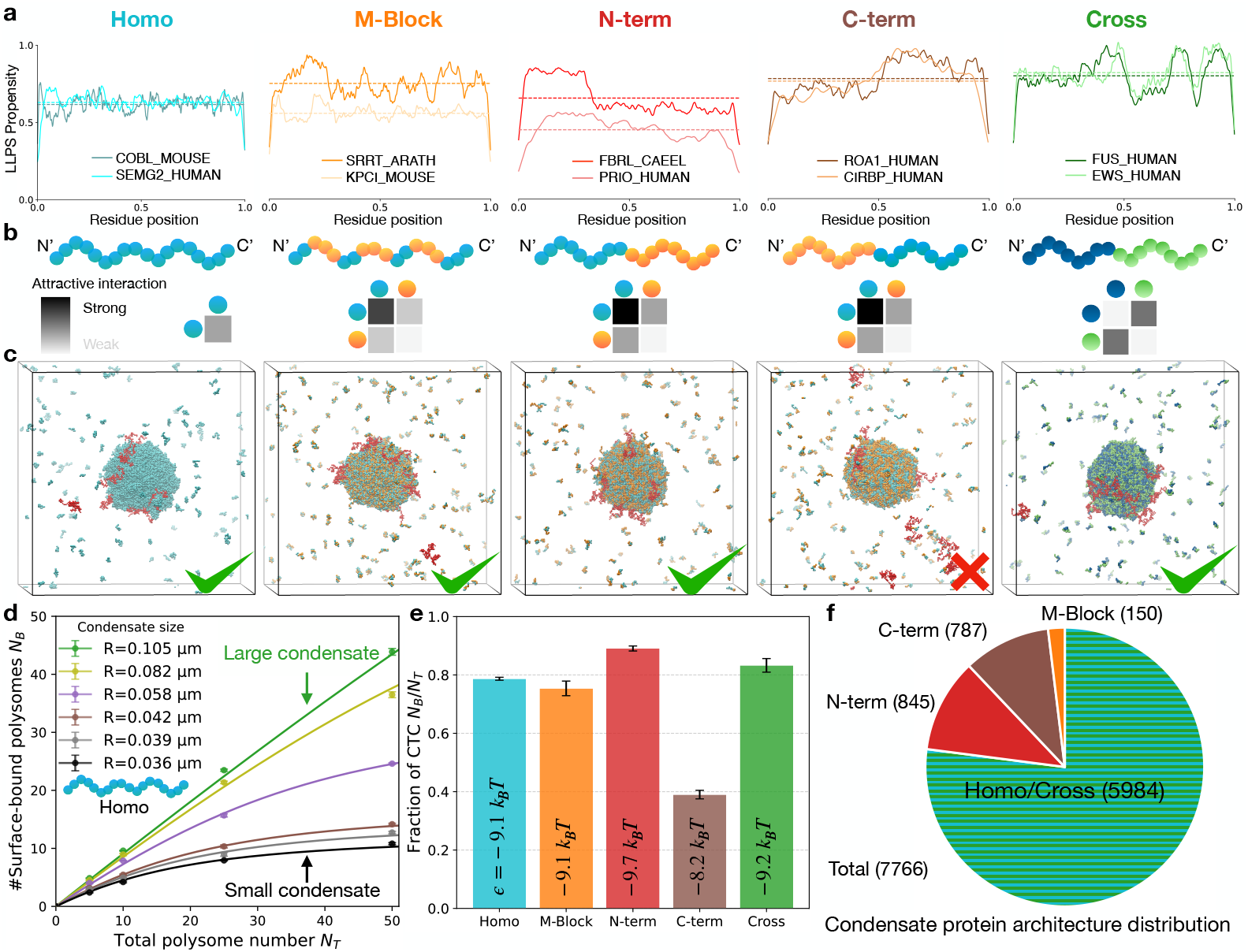
Protein architecture influences the extent of Co-Translational Condensation. **a**, Predicted LLPS propensity profiles along the sequence of representative proteins from five distinct architecture classes, see Methods. Regions exceeding the average LLPS propensity (dashed line) indicate strong-interaction regions for that protein. We note that the Cross examples are annotated by experimental results [28] because based on LLPS propensity we cannot distinguish between Homo and Cross architectures. **b**, Schematics for the architectures with interaction legends shown below; darker shading indicates stronger attraction between regions. **c**, Simulation snapshots showing the spatial distribution of polysomes and proteins for each architecture. All classes except C-term show substantial CTC. See SI Movies II-VI for full trajectories. **d**, Surface-bound polysome number *N*_*B*_ versus total polysome number *N*_*T*_ (Homo architecture, see Extended Data Fig. 2 for others). Data points represent mean values from 3 independent simulations, where the error bars show the standard error. Different colored points show different condensate sizes. The curves show the global fit to a Langmuir adsorption model Eq. 2. *N*_*B*_ plateaus for small condensates because the surface becomes saturated. The number increases linearly for large condensates, reflecting an unsaturated surface. Data shown for **c-e** are results from small polysomes, see Extended Data Fig. 2 for all other setups. **e**, Fraction of surface-bound polysome (fraction of CTC *f* = *N*_*B*_*/N*_*T*_) versus total polysome number *N*_*T*_ for a medium-sized condensate (*R* = 0.062 *µ*m). The curves show the global fits for each architecture. N-term and Cross architecture show strong CTC and C-term shows the weakest. **f**, Distribution of protein architectures in the CD-CODE database (N=7766) [27]. For details of the classification and additional analysis, see Methods.

- **Homo**: Interaction regions are evenly distributed along the sequence.
- **M-Block**: Multiple strong-interaction blocks alternate with weak ones.
- **N-term**: Strong-interaction regions are concentrated near the N-terminus.
- **C-term**: Strong-Interaction regions are concentrated near the C-terminus.
- **Cross**: The N- and C-terminal regions preferentially interact with each other, rather than with themselves.

To compare their CTC behavior, we constructed coarse-grained bead-spring models for each architecture (Fig. 2**b**). Interaction strengths were calibrated to yield approximately equal saturation concentrations across all classes, ensuring that observed differences in CTC stem from architecture rather than phase-separation driving forces (see Extended Data Fig. 1, 3). Notably, N-term and C-term are mirror images; while indistinguishable for phase separation, they differ in which part of the protein is displayed in the nascent chains of their polysome.

Fig. 2**c** shows that different architectures lead to different degrees of CTC. We quantified this by fitting the number of co-localized (surface-bound) polysomes *N*_*B*_ to a Langmuir adsorption model on a spherical surface with a finite reservoir and an ideal-chain unbound state (see Methods). The corresponding free energy in units of *k*_*B*_*T* is

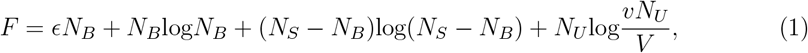

where *N*_*B*_ is the number of bound polysomes, *ϵ* is the corresponding binding energy, *N*_*S*_ is the maximum number of polysomes that can fit on a condensate surface (*N*_*S*_ = 4*πR*^2^*/σ*) with condensate radius *R*, and *σ* is the polysome binding area. *N*_*U*_ is the number of unbound polysomes in the dilute phase, *v* is the volume of a polysome, and *V* is the dilute-phase volume (finite reservoir). Note *N*_*T*_ = *N*_*B*_ + *N*_*U*_ = const.

At equilibrium, minimizing the free energy with respect to *N*_*B*_ yields

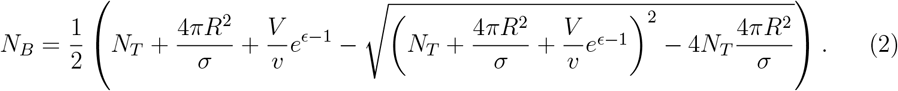

For each protein architecture, we globally fit *N*_*B*_ from simulations across different *N*_*T*_ and *R* to extract *ϵ* and *σ* as characteristic parameters. The adsorption model agrees well with simulations for all architectures (Fig. 2**d**, Extended Data Fig. 2). For small condensates, *N*_*B*_ plateaus at a moderate *N*_*T*_, indicating a saturated surface In contrast, larger condensates exhibiting more binding sites, favoring CTC. Notably, large condensates are more representative of the cellular environment, where the condensate surface is far from saturation limit and *ϵ* primarily governs the extent of CTC.

Therefore, we quantify the CTC ability of each architecture via *ϵ* in Eq. 3.1 (Fig. 2**e**). Among the five architectures, N-term and Cross exhibits the strongest CTC, as all the strong interaction regions are fully exposed at tips of the polysome branches. In contrast, C-term shows weak CTC because the strong interaction regions are only exhibited by a few nearly completed nascent proteins.

Finally, Fig. 2**f** shows that most condensate-associated proteins (79%) exhibit Homo, Cross, or M-Block architectures, all comparably favorable for CTC. The C-term class, which exhibits weak co-localization, accounts for only 10%. The 11% share of N-term proteins represent the strongest candidates for CTC (see SI for list of N-term proteins), further signifying the prevalence of architectures that favor CTC.

### 3.2 Large polysome stabilize small condensate

Polysomes have more protein interaction regions through their multiple nascent chains than a single protein, giving them a disproportionate influence on the phase diagram. This effect is amplified in large polysomes with more nascent chains. However, the fraction of proteins present as nascent chains in polysomes is less than 0.1% of the total protein pool[24]. We therefore asked whether this disproportionate influence can overcome their marginal concentration.

The impact of polysomes on the phase diagram is quantified by how *c*^Dilute^ changes with increasing polysome number. As shown in Figs. 3**a**,**b**, small polysomes do not alter *c*^Dilute^ for condensates of any size. In contrast, large polysomes promote phase separation (lowering *c*^Dilute^) for small condensates. Large condensates are unaffected. We note that *c*^Dilute^ is condensate–size dependent due to surface tension[5], distinguishing it from the binodal of an infinite system. All architectures behave similarly; see Extended Data Fig. 3 for further discussion. Since biologically relevant condensates are typically larger than 100 nm, our results indicate that binodals measured experimentally reflect the underlying protein phase behavior and can safely be interpreted without considering polysomes.

**Fig. 3:**
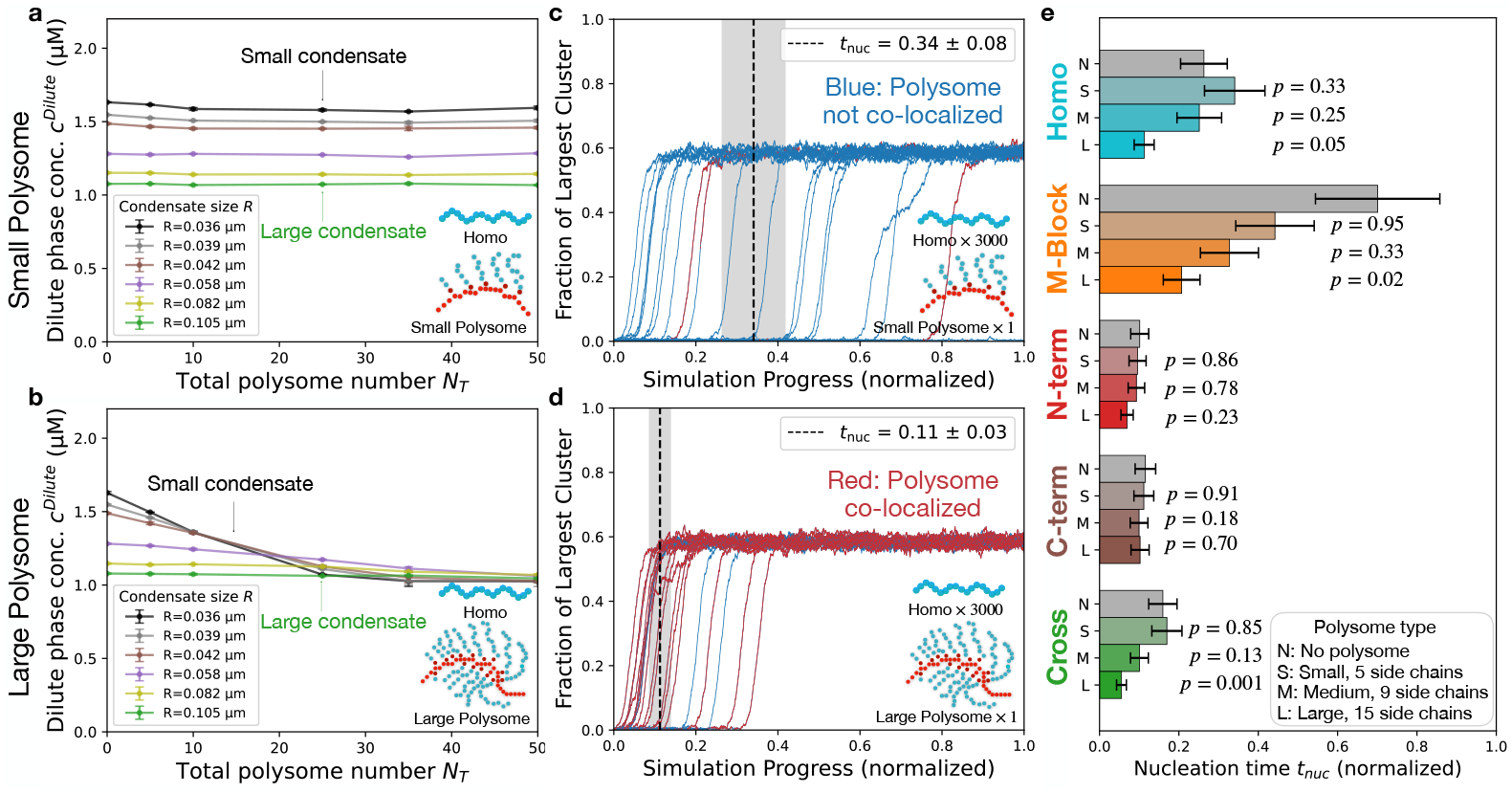
Large polysomes impact phase behavior and serve as nucleation centers. Dilute phase concentration *c*^Dilute^ versus numbers of polysomes *N*_*T*_ for different average condensate sizes *R*. Error bars denote standard errors. Curves are guides to the eye. Data shown is for Homo, see Extended Data Fig. 3,4 for other architectures and polysome sizes. **a**, Small polysomes have negligible impact on condensate thermodynamics. **b**, Large polysomes perturb the thermodynamics of small condensates (black) significantly but not large condensates (green). For discussion on surface tension effects, see SI. **c**,**d** Fraction of proteins in the largest cluster (condensate) with 3000 proteins and one polysome. Each curve corresponds to an independent simulation (20 replicas). **c**: Small polysome; **d**: Large polysome. The nucleation time is defined as the last point before the largest cluster reaches 5% of total proteins, and the plateau at ∼ 60% corresponds to the equilibrium condensate. Red/blue indicate co-localization/de-localization with the cluster (shown for Homo; see SI for simulations without polysomes and with other architectures). Dashed lines mark the mean nucleation time, with standard error in gray. These results show that large polysomes accelerate condensate nucleation by serving as nucleation centers. **e**, Summary of the nucleation time for different protein architectures with error bars indicating standard errors. The statistical significance p-values of different polysome sizes (red) by comparing to no polysome case (gray) are labeled, *p <* 0.05 means the two cases are statistically significantly different.

## 4 Kinetics of Co-Translational Condensation

Polysomes have finite lifetimes, raising the question of whether CTC, even if thermodynamically favored, can occur quickly enough. We examine if this can happen through two routes: either condensates nucleate directly on polysomes, or polysomes diffuse to existing condensates.

### 4.1 Large polysomes nucleate new condensates

To investigate nucleation on polysomes, we simulated condensate formation with one polysome, see Methods. Fig. 3**c,d** shows the size of the largest cluster as a function of time for 20 independent simulations. Random fluctuations lead to the formation of a small cluster which grows into an equilibrated condensate (∼ 60% of the total proteins). The polysome may (red) or may not (blue) be co-localized with the nucleation and growth of condensates. While small polysomes are scarcely co-localized with a growing condensate, see Fig. 3**c**, large polysomes display a strong correlation with nucleation of condensates.

We define the nucleation time ⟨*t*_nuc_⟩ as the average of the times when individual simulations cross a cluster size threshold (5%), see Methods. We quantify the impact from polysomes by changes in ⟨*t*_nuc_⟩ compared to simulations without a polysome ⟨*t*_nuc_⟩ = 0.26 *±* 0.06, see Extended Data Fig. 4.

A large polysome reduces ⟨*t*_nuc_⟩ by a factor of three (0.11 *±* 0.03, *p* = 0.052). In contrast, the effect of a small polysome on ⟨*t*_nuc_⟩ is not statistically significant (0.34 *±* 0.08, *p* = 0.334). This is consistent with our observation that large polysomes co-localize with nucleation events, whereas small polysomes do not. Moreover, the formation of new condensates is limited by the stability of small clusters. As shown in Fig. 3**a,b**, large polysomes stabilize condensates in a size-dependent manner, with the effect strongest for the smallest condensates, whereas small polysomes show no measurable stabilization.

Fig. 3**e** shows the impact of the architectures on nucleation. Enhanced nucleation from large polysomes are observed only for Homo, M-Block, and Cross architectures but not for N-term or C-term.

Large polysomes promote nucleating a condensate on each polysome once the saturation concentration is exceeded. By stabilizing otherwise unstable small condensates, they effectively lower the nucleation barrier and accelerate condensate formation. In addition, de-localized large polysomes generate a locally supersaturated environment through translation, further driving proteins to nucleate into condensates on themselves.

### 4.2 Polysomes diffuse to existing condensates

Polysomes can also reach condensates via diffusion. The fraction of CTC is limited by both the thermodynamics of adsorption and the ratio between the diffusion and disassembly time scales. Multiple condensates usually coexist within a single cell – alleviating the impact of diffusion length scales[29] by decreasing the average distance between condensates *L*, see Fig. 4**a**. Multiple condensates can be taken into account by considering one condensate and using reflective boundary conditions at the distance *L/*2.

**Fig. 4:**
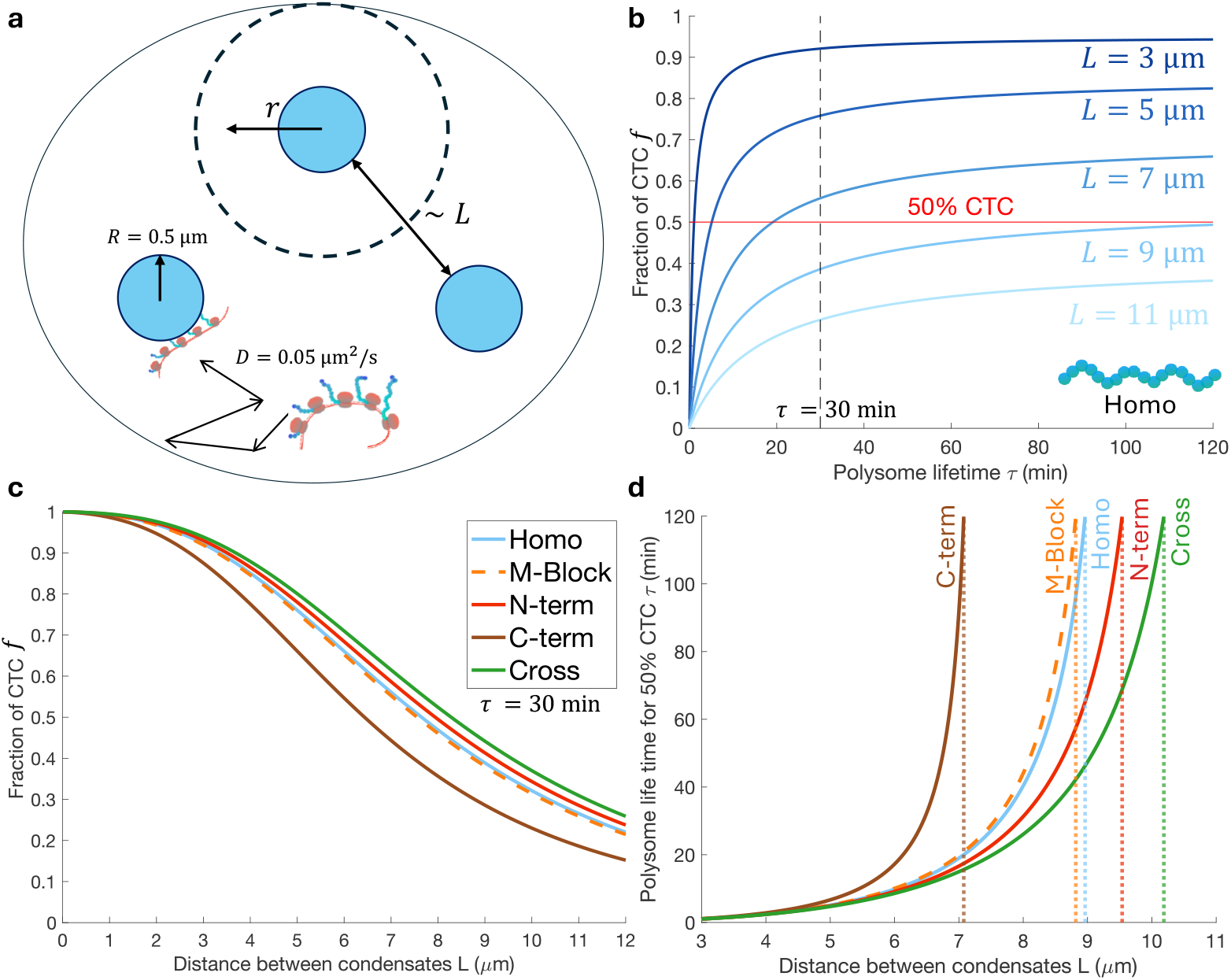
Combined dynamics and thermodynamics of Co-Translational Condensation. **a**, Cartoon of reaction-diffusion model. Typically, a cell contains multiple condensates of the same kind, where *L* describes the average distance between condensates. This reduces the diffusion length scale so we model from a condensate to the mid-point between neighboring condensates with a spherical symmetry. The thermodynamics of binding at the condensate surface are chosen to match the Langmuir model Eq. 4 and see SI for details. **b**, Fraction of CTC *f* versus polysome life time *τ*. Different lines denotes different distances between condensates *L*. This suggests that for a moderate spacing (*L* ∼ 6 *µ*m), a medium-life-time polysome (*τ* ∼ 30 min) can reach 50% CTC. **c**, Fraction of CTC *f* versus distance *L* for *τ* = 30 min for all architectures (different colors). For small *L*, most polysomes are bound to the surface. For medium *L, f* is limited by the thermodynamics of the architecture. For large *L, f* is limited by the kinetics. Binding energy used here is extracted from the medium polysome setup, see Methods for details. **d**, Polysome lifetime required for 50% CTC. There is a maximal value of *L* beyond which 50% CTC is no longer possible. This is set by the thermodynamics of binding, hence each architectures has its own maximal value of *L*^∗^ (dotted lines, *τ* = 120 min).

Diffusion and disassembly of polysomes are modeled by a reaction-diffusion equation with spherical symmetry for the region outside of the condensate of radius *R*:

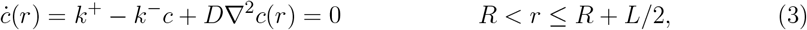

where *c*(*r*) is the polysome concentration at radius *r, k*^+^ is the polysome assembly rate, and *k*^−^ is the disassembly rates defined by the polysome’s life time *τ* = 1*/k*^−^, respectively, and *D* = 0.05 *µ*m^2^*/*s is the diffusion coefficient of the polysome.

At the condensate surface *R* = 0.5 *µ*m, we imposed an absorbing boundary condition governed by the Langmuir adsorption model (Eq. 3.1, Fig. 2**d,e**),.

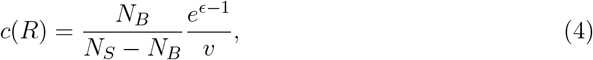

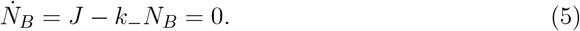

Here, *ϵ* is the binding energy obtained from from the coarse-grained simulation for each architecture with medium polysomes, see Methods. The incoming flux 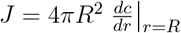 and polysomes bound (*N*_*B*_) to the condensate also disassemble at the rate *k*^−^.

As discussed above, a reflective boundary condition is imposed at the mid-point between condensates *R*_*D*_ = *R* + *L/*2,

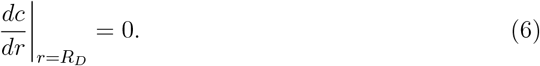

If only one condensate exists then *R*_*D*_ is the radius of the cell.

The resulting fraction of CTC is defined as the fraction of polysomes bound to the surface *f* = *N*_*B*_*/N*_*T*_, see Methods.

In Fig. 4**b**, we show how the fraction of CTC depends on polysome lifetime *τ* = 1*/k*^−^ and distance between condensates *L*. When condensates are close together, even shortlived polysomes achieve high CTC (*L* = 3 *µ*m, *τ* = 5 min → *f* = 70%). At intermediate spacing, a longer lifetime is required to reach *f* = 50% (*L* = 6 *µ*m, *τ* = 30 min). At large separations, binding of polysomes to condensates is unlikely such that there is no lifetime long enough to achieve a high fraction of CTC (*L* = 9 *µ*m, *τ* = 120 min → *f* = 30%).

Due to differences in binding energies *ϵ*, different architectures yield varying levels of CTC at a fixed lifetime. Here we chose *τ* = 30 min as a reference for a typical lifetime of polysomes in yeast[30, 31]. Fig. 4**c** shows how *f* depends on the separation between condensates for different architectures. As expected from the analysis of effective adsorption energy *ϵ* (Fig. 2**e**), N-term produces the strongest CTC even for moderate *L*, while C-term suppresses CTC for all but the shortest distances between condensates.

To quantify the max separation between condensates where CTC can occur, we define the critical distance *L*^∗^ beyond which 50% CTC is no longer achievable within a polysome lifetime of *τ* = 120 min. As shown in Fig. 4**d**, each sequence has a distinct *L*^∗^, beyond which the required lifetime diverges. For example, N-term maintains efficient CTC up to *L*^∗^ ≈ 8 *µ*m, whereas C-term loses CTC potential beyond *L*^∗^ ≈ 4 *µ*m.

CTC is expected to occur through diffusion of polysome to condensate surfaces with moderate separation between condensates and average polysome lifetimes. The condensate separations that are conducive to CTC are seen in eukaryotic cells[1, 32, 33] and are the sizes of small organisms such as yeast[10]. We note that small polysomes can diffuse faster to existing condensates than large polysomes, allowing CTC with shorter lifetime of polysomes.

## 5 Functional implications of Co-Translational Condensation

### 5.1 CTC facilitates efficient Post-Translational Modifications

Biomolecular condensates are active reaction crucibles that can spatially organize and regulate biochemical processes. One potential functional benefit of CTC is the enhanced efficiency of protein Post-Translational Modifications (PTMs).[2, 34–38].

To illustrate this, we consider a simple reaction in which an immature protein species *A* is converted into its functional form, species B, via an enzymatic reaction localized within the condensate (Fig. 5**a**). The reaction could represent phosphorylation, SUMOylation, or other enzyme-mediated PTMs.

**Fig. 5:**
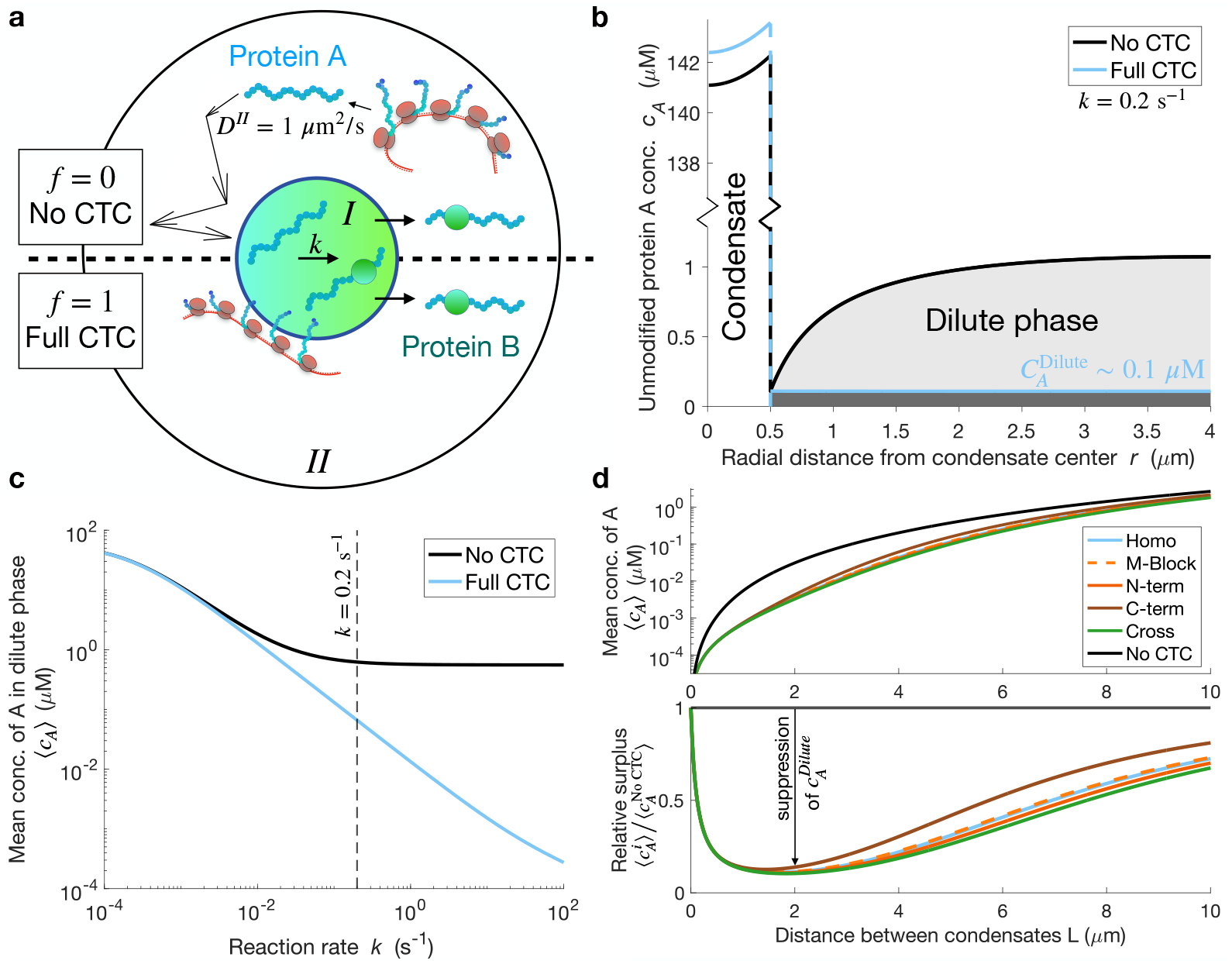
CTC facilitates protein post-translational modifications. **a**, Model system of post-translational protein modifications in condensates, *A* → *B*. Top half *f* = 0: Translation occurs without CTC – the unmodified protein *A* has to diffuse in the cytoplasm to locate the condensate before it can be modified. Bottom half *f* = 1: Translation occurs with CTC – protein *A* is synthesized directly at the surface of the condensate and is immediately ready for modification. **b**, Concentration profile of unmodified protein *A* for two cases. With no CTC *f* = 0 (black), the concentration profile shows a large accumulation of material that is diffusing to the condensate. We note the break in the y-axis to accommodate the visualization of the condensate. With full CTC *f* = 1 (blue), the concentration profile outside the condensate is low and flat. Solutions correspond to condensates of size *R* = 0.5 *µ*m and distances between condensates of *L* = 7 *µ*m. **c**, The mean concentration of *A* in the cytoplasm as a function of PTM reaction rate. When the reaction rate is higher, the benefit of having CTC is more pronounced because diffusion dominates the behavior of the no CTC *f* = 0 condition (black) while CTC *f* = 1 shows little diffusion dependence (blue). **d**, Unifying the PTM dynamics with the combined dynamics and thermodynamics of CTC. Top: Mean concentration of *A*, no CTC *f* = 0 (black) and all five architectures 0 *< f*^*i*^(*L*) *<* 1 (colored), where *f*^*i*^(*L*) is the emerging fraction of CTC for each architecture *i* and distance between condensates *L* (Fig. 4 **c**, medium polysomes). Bottom: Degree of impact from CTC with different architectures. The relative surplus is defined as the mean concentration in the cytoplasm with different architectures compared to the null case of no CTC. The suppression of *A* is the distance from unity. Architectures with greater CTC propensity achieve stronger suppression and maintain this effect across a broader range of condensate spacings.

In the absence of CTC (*f* = 0), illustrated in Fig. 5**a** upper half, protein *A* is synthesized and degraded throughout the cytoplasm (Phase *II*: *R < r < R*_*D*_) and is converted into *B* inside a condensate (Phase *I*: 0 *< r < R*) which must be reached via diffusion. With full CTC (*f* = 1), illustrated in Fig. 5**a** lower half, polysomes are co-localized, such that synthesis of protein *A* occurs directly at the condensate surface. The reactions and diffusion at steady-state are described by

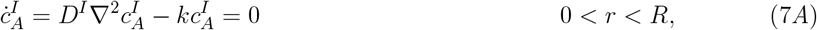

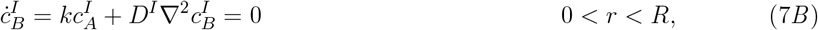

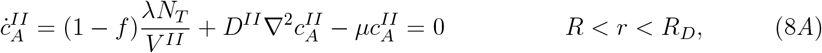

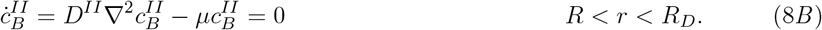

Here, *c* is the radial protein concentration with superscripts *I/II* denoting the dense/dilute phases respectively and subscripts *A/B* denoting unmodified/PTM protein species, *D* is the diffusion coefficient in the respective phase, *k* is the PTM reaction rate, *µ* is the protein degradation rate, *f* is the fraction of CTC as discussed above, and *λN*_*T*_ is the total translation rate of the protein *A*.

At the condensate interface, we impose a protein partition factor of *P* = 1300, which is extracted from our simulations and is consistent with experimental measurements[39]:

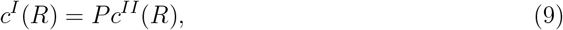

and a flux balance condition

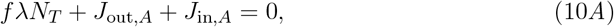

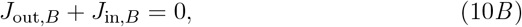

where *fλN*_*T*_ is the synthesis at the surface, 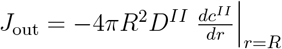 is the outward flow just outside the condensate, and 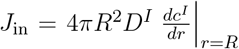 is the inward flow just inside the condensate. Additionally, we impose regular behavior at the center of the condensate 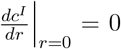 and, as before, reflective boundary condition at the mid-point between condensates 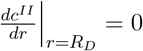.

First, we compare the amount of unmodified protein *A* in two limiting cases of no CTC (*f* = 0) and full CTC (*f* = 1). Fig. 5**b** shows the concentration profiles for both limits: less unmodified protein *A* accumulates in the cytoplasm with CTC compared to no CTC. In Fig. 5**c** we show the mean concentration of *A* averaged over the diluted phase as a function of the PTM reaction rate *k*. From this we can conclude that the efficiency increase of PTM by CTC is significantly stronger when the enzymatic conversion of *A* to *B* is fast compared to diffusion (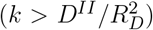). The accumulation of unmodified protein *A* is inefficient from a resource allocation perspective and may be undesirable if protein *A* is aggregation-prone or toxic. With CTC, by localizing translation to the condensate surface, protein *A* is born in the condensate and can be promptly modified.

Next, we considered the intermediate case where the adsorption and diffusion control the fraction of CTC (0 *< f <* 1). Specifically, we used the fraction of CTC previously calculated in the polysome diffusion model (Fig. 4**c**) to find the mean concentration of *A* in the dilute phase ⟨*c*_*A*_⟩. Fig. 5**d** top shows ⟨*c*_*A*_⟩ as a function of the distance between condensates *L*. As condensates are more sparsely distributed in cells (increasing *L*), ⟨*c*_*A*_⟩ rises.

Importantly for CTC consideration is how much having CTC changes things compared to no CTC; this motivates us to examine the relative surplus of *A*, 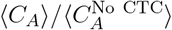. Fig. 5**d** bottom shows the surplus as a function of distance between condensates. Dilute phase *A* is maximally suppressed 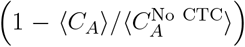 for intermediate condensate spacing, see SI.

Architectures with greater CTC propensity achieve stronger suppression of dilute phase *A* and maintain this effect across a broader range of condensate spacings. In other words, the easier it is for a given protein architecture to achieve CTC, the more effectively it reduces the accumulation of unmodified protein *A* in the cytoplasm – even as condensates become more sparsely distributed. This highlights how protein design not only determines whether CTC occurs, but also how robustly it functions across varying cellular conditions.

## 6 Outlook: Interpreting Protein Organization and Evolution

Our study establishes Co-Translational Condensation (CTC) as a general mechanism that links translation to phase separation. The central feature is that polysomes, by virtue of their many nascent chains, can adsorb onto condensates. Using coarse-grained simulations, we demonstrate that protein domain architecture sets the strength of this adsorption, and bioinformatic analysis suggests that most condensate-associated proteins have architectures favorable for CTC. These findings are consistent with recent experiments showing that translating mRNAs co-localize with the corresponding condensates [10, 11].

Using multi-scale modeling, we show polysomes can associate with condensates either by diffusion or by acting as nucleation centers. In particular, large polysomes lower the nucleation barrier, promoting the formation of multiple condensates of one type, as observed in cells.

CTC could offer physiological advantages. Often enzymes modify proteins to provide them with the desired biological function after translation. We have demonstrated that CTC can accelerate such post-translational modifications, thereby minimizing intermediates and enhancing signaling responsiveness. Additionally, CTC could lead to co-translational modifications (modifications that occur during translation). CTC may be especially important in pathways requiring rapid feedback, such as immune response or stress dynamics.

Co-translational folding could also be impacted by CTC for good or bad. Folded protein structures depend on the local chemical environment [40, 41], the presence of chaperones [42], and translational pauses [43]. Co-translational folding has been shown to assist all three of these[44]. CTC could influence each of these aspects as well by bring translation to condensates that have distinct chemical environments[45, 46], concentrate chaperones [47, 48], and alter the local pool of tRNAs [49].

A previous study has focused on proteins that form homo-oligomers with themselves. They showed that the oligomerization-regions of the proteins are preferentially located at the C-terminal, presumably to avoid premature self-association and aberrant aggregation during translation [50]. Our bioinformatic analysis suggests that proteins that are associated with condensates shows the opposite trend where architectures favor co-translational interactions. This suggests different evolutionary pressures between oligomerization-proteins and condensate-proteins. Additionally, similar proteins can belong to different architecture classes. An example of this is the FET family of proteins, where the prion-like domains can be located at either terminus[28].

We propose that CTC represents a poorly explored axis along which protein function, folding fidelity, and spatial organization could be regulated. Our coarse-grained architectures offer a conceptual framework for predicting which proteins are good candidates for CTC. This framework may also prove useful for informing the rational design of synthetic systems – either to harness CTC for enhanced efficiency or to avoid it when premature interactions are detrimental.

## Supplementary information

The Supporting Information is available:

Simulation methodology, design, and convergence, diffusion-reaction models, and video captions (PDF)

## Acknowledgements

We thank Emmanuel Levy and Meta Heidenreich for providing the experimental data used for Fig. 1**d** and insightful discussions. We are grateful to Patrick McCall, Xi Chen, Simon Alberti, Titus Franzmann, Arash Nikoubashnman, and Louise Jawerth and for feedback and discussion. J.U.S. acknowledges support from the Deutsche Forschungsgemeinschaft (DFG) under the grant number SO-277/25. The authors acknowledge the Center for High-Performance Computing (ZIH) Dresden for computational resources.

## Declarations

Some journals require declarations to be submitted in a standardised format. Please check the Instructions for Authors of the journal to which you are submitting to see if you need to complete this section. If yes, your manuscript must contain the following sections under the heading ‘Declarations’:

- Funding
- Conflict of interest/Competing interests (check journal-specific guidelines for which heading to use)
- Ethics approval and consent to participate
- Consent for publication
- Data availability
- Materials availability
- Code availability
- Author contribution

If any of the sections are not relevant to your manuscript, please include the heading and write ‘Not applicable’ for that section.

